# Monitoring Sphingomyelin Biosynthesis at Nanoscale Resolution by Expansion Microscopy

**DOI:** 10.64898/2026.05.27.726241

**Authors:** Fabienne Wagner, Arpita Mohanty, Fabian Schumacher, Julian Fink, Louise Kersting, Burkhard Kleuser, Jürgen Seibel, Vera Kozjak-Pavlovic, Thomas Rudel, Marcel Rühling

**Affiliations:** Chair of Microbiology, Julius-Maximilians-University Würzburg, Theodor-Boveri-Weg, 97074 Würzburg, Germany; Institute of Pharmacy, Freie Universität Berlin, Königin-Luise-Straße 2+4, 14195 Berlin, Germany; Institute of Organic Chemistry, Julius-Maximilians-University Würzburg, Am Hubland, 97074 Würzburg, Germany

## Abstract

Sphingomyelin is the most abundant sphingolipid in mammalian cells and is synthesized by two isoenzymes, sphingomyelin synthase 1 and 2, located in the Golgi and in the plasma membrane. Abnormal sphingomyelin synthesis is associated with infections and diseases such as diabetes and cancer. Measuring cellular sphingomyelin synthase activity fosters our understanding of how these enzymes are involved in pathological processes and can be crucial for the identification of therapeutic compounds modifying sphingomyelin biosynthesis. We have developed a novel fluorometric assay that enables microscopic detection of cellular sphingomyelin synthase activity. We show that sphingomyelin synthases use propargyl choline and α-NH_2_-ω-N_3_-C_6_-ceramide to generate trifunctional sphingomyelin, a lipid derivative detectable with spatial resolution via Förster resonance energy transfer. By combining this assay with expansion microscopy, a super-resolution imaging technique, we measured the distribution of *de novo* synthesized trifunctional sphingomyelin in cells at nanoscale resolution, thereby directly demonstrating sphingomyelin biosynthesis at the Golgi and the plasma membrane. By monitoring sphingomyelin biosynthesis and degradation in cells infected with the obligate intracellular pathogen *Chlamydia trachomatis*, we dissected the complex sphingolipid metabolization of these bacteria with unprecedented resolution. By correlating spatial metabolic information with lipidomics, we provide a powerful tool for investigating cellular sphingomyelin metabolism.

**Teaser:** Combining functional lipids, expansion microscopy and FRET helps visualise sphingolipid metabolism at unprecedented resolution.

## INTRODUCTION

Sphingolipids are not only major constituents of eukaryotic membranes but also possess important functions as bioactive signaling molecules [1–4]. Sphingomyelin (SM), the most abundant sphingolipid in mammalian cells, is thought to be an important component of lipid microdomains (also called lipid rafts), which were often described to influence cellular signaling processes for instance by affecting the distribution of proteinaceous receptors on the plasma membrane [5–9].

Ceramide (Cer), a central molecule in sphingolipid metabolism, is synthesized from serine and palmitoyl-CoA over multiple steps in the endoplasmic reticulum (ER) [10]. Cer can be further modified to yield so-called complex sphingolipids, for example by addition of carbohydrate residues to form glycosphingolipids and gangliosides or by attachment of a phosphocholine head group resulting in the formation of SM [10, 11]. For SM biosynthesis, Cer is shuttled from the ER to the Golgi apparatus either by vesicular transport or via the Cer transfer protein (CERT) [10, 12]. Within the Golgi, two isoenzymes, sphingomyelin synthase 1 and 2 (SMS1 and SMS2, respectively), catalyze the synthesis of SM by transferring a phosphocholine residue from phosphatidyl choline (PC) to Cer, thereby releasing diacylglycerol (DAG) [13–18]. While SMS1 is exclusively found in the Golgi, SMS2 additionally localizes to the plasma membrane [16–18], where it regulates SM levels and influences signaling processes [19, 20]. SMS deficiency is accompanied by changes in cellular sphingolipid profiles, thereby not only decreasing SM levels but also resulting in the accumulation of other lipid species such as glycosphingolipids [21, 22] or Cer [23].

Stimulating or interfering with SM biosynthesis has been suggested as a therapeutic approach. For instance, antitumor medication has been demonstrated to restore abnormalities in the lipid profile of cancer cells by increasing SMS activity [24]. By contrast, inhibition of SMS2 has been proposed as a medication for breast cancer [20], type 2 diabetes [25], obesity [26], and atherosclerosis [27]. SM biosynthesis also fosters infection by a variety of viruses, including, rubella virus [28], Japanese encephalitis virus [29], influenza virus [30], and severe fever with thrombocytopenia syndrome virus [31].

The obligate intracellular pathogen *Chlamydia trachomatis* is a leading cause of sexually transmitted diseases [32]. The bacteria are predominantly found in two developmental forms, the smaller infectious elementary bodies (EBs) and the metabolically active and replicative reticulate bodies (RBs) [33, 34]. EBs adhere to and are taken up by the host cell in a membrane-bound compartment and subsequently undergo a differentiation process to form RBs [35–38]. Thereafter, *C. trachomatis* forms a large vacuole within the host cell, the so-called inclusion, which serves as replicative niche [33, 34]. Sphingolipids are crucial for *C. trachomatis* infection as they facilitate inclusion stability and bacterial replication [39, 40]. Due to a very reduced genome comprised of only ~900 genes [41], *C. trachomatis* is highly dependent on host cell metabolites, including lipids [42]. *C. trachomatis* acquires sphingolipids from the host cell either by redirecting vesicles from the Golgi to the inclusion [43] or by the recruitment of CERT to the inclusion membrane [42, 44–47]. Subsequently, the lipids are incorporated into bacterial membranes [42, 44–47]. Despite the general metabolic dependence on the host, previous studies suggested that *C. trachomatis* encodes an enzyme that synthesizes SM [48, 49], highlighting the significance of this molecule for the infection.

Identification of novel compounds affecting *de novo* SM synthesis requires reliable assays to monitor cellular SMS activity. Previously, SMS activity was measured by incubating cells with fluorescently-tagged Cer analogs, such as NBD-C_6_-Cer, which is converted to the corresponding SM derivative. Subsequently, lipids were extracted, and the proportion of *de novo* synthesized SM was determined either by thin-layer chromatography (TLC) [50–52] or by liquid chromatography-tandem mass spectrometry (LC-MS/MS) [53]. We previously used functionalized SM derivatives, so-called trifunctional sphingomyelin 1 and 2 (TFSM1 and TFSM2, respectively), to visualize cellular SM distribution and to monitor SM degradation during *C. trachomatis* infection via ExM [45, 60, 61] α-NH_2_-ω-N_3_-C_6_-Cer [44] and propargyl-choline (ppCho, [62]) by human cells, which can be quantified by Förster resonance energy transfer (FRET). To distinguish between synthetic TFSM2 that can be used to follow SM catabolism, we term TFSM2 generated by living cells “*bio*TFSM2”. Using liquid chromatography-tandem mass spectrometry (LC-MS/MS) and FRET acceptor bleaching, we demonstrate that *bio*TFSM2 is synthesized by human cells from the two artificial substrates. TFSM generation was only observed in HeLa WT cells, while cells deficient in SMS1 and 2 (SMS dK.O., [63]) could not generate *bio*TFSM2. Combining FRET acceptor bleaching with ExM enables the measurement of *bio*TFSM2 content with subcellular resolution using conventional confocal microscopes. For the first time, we show by microscopy that the Golgi apparatus and the plasma membrane contain higher levels of *de novo* synthesized SM compared to other cellular compartments such as mitochondria and the nuclear envelope. In addition, we monitored *bio*TFSM *de novo* synthesis and TFSM degradation during *C. trachomatis* infections via ExM. By correlating the spatial information with lipidomics, we dissected the complex sphingolipid metabolism in *C. trachomatis* inclusions with unprecedented details at single-bacterium level. Our assay represents, therefore, a powerful and accessible tool for visualization and quantification of SMS activity and SM synthesis in both human cells and bacteria.

## RESULTS

### Design of the SMS assay

(*Bio)*TFSM2 possesses three functional groups. The primary amino function (**Fig. 1**, orange circle) enables fixation of the molecule with aldehyde fixatives and is indispensable for embedding the compound into the hydrogel during ExM [44, 45]. Furthermore, there are two functional groups that can be targeted by click chemistry: an alkyne group (blue circle) in the phosphocholine head and an azide function (magenta circle) in the acyl side chain. The azide function can be equipped with a fluorophore via strain-promoted azide-alkyne cycloaddition (SPAAC) without the requirement for additional catalysts, e.g. by using dibenzocyclooctyne (DBCO)-coupled dyes. By contrast, mounting a fluorophore to the alkyne function of (*bio*)TFSM2 requires copper-catalyzed azide-alkyne cycloaddition (CuAAC), thereby facilitating a controlled and selective attachment of fluorescent dyes to (*bio*)TFSM2.

**Fig. 1.**
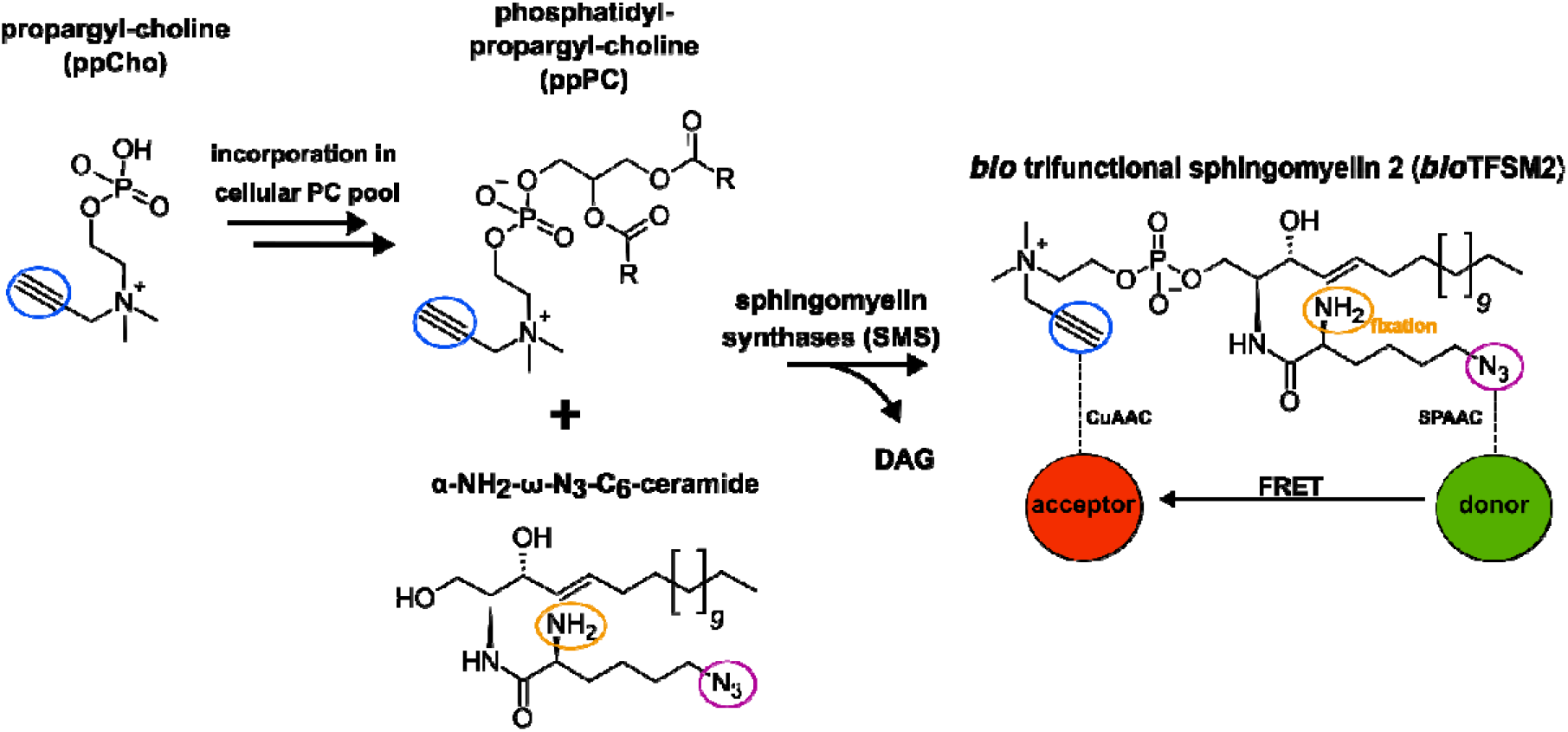
Generation of *bio*TFSM2 by SMS from artificial substrates. Treatment of cells with propargyl choline (ppCho) results in incorporation of ppCho into the cellular phosphatidyl choline (PC) pool yielding in phosphatidyl-propargyl-choline (ppPC). ppPC and α-NH_2_-ω-N_3_-C_6_-ceramide are converted to *bio* trifunctional sphingomyelin 2 (*bio*TFSM2) and diacylglycerol (DAG) by SMS. The primary amine function (orange circle) enables the fixation of *bio*TFSM2 by aldehyde fixatives. *bio*TFSM2 can be equipped with a FRET pair by attaching fluorophores to the azido group (magenta circle) via strain-promoted azide-alkyne click chemistry (SPAAC) and to the alkyne function (blue circle) via copper-catalyzed azide-alkyne click chemistry (CuAAC).

We previously showed that TFSM2 is converted to the respective Cer and sphingosine metabolites by human cells [45, 61]. If the two fluorophores used for click chemistry constitute a FRET pair, the proportion of non-metabolized native TFSM2 correlates with the intensity of the detected FRET signal, which can be used to measure TFSM2 degradation.

Here, we aimed to monitor SM *de novo* synthesis by monitoring the generation of *bio*TFSM2 within living cells. Therefore, we used ppCho [62] and α-NH_2_-ω-N_3_-C_6_-Cer [44], which in our assay serve as artificial substrates for *bio*TFSM2 synthesis. ppCho was shown to be incorporated efficiently into the PC pool of mammalian cells resulting in phosphatidyl-propargyl-choline (ppPC) [62].

ppPC is likely accepted by at least one of the two SMSs as suggested by the incorporation of ppCho in SM species observed in earlier studies [62, 64]. In contrast to TFSM2, α-NH_2_-ω-N_3_-C_6_-ceramide and their metabolites, which contain a primary amino function, many natural lipids are not retained in samples fixed with aldehyde fixatives upon permeabilization with harsh detergents such as Triton X-100 [44, 45, 65]. Accordingly, this results in removal of most other ppCho-labeled lipids, including ppPC, thereby minimizing fluorescence that does not originate from TFSM2 and its metabolites.

Hence, we hypothesized that cellular SM biosynthesis can be monitored through the generation of *bio*TFSM2 from ppCho and α-NH_2_-ω-N_3_-C_6_-Cer and thus, cellular SMS activity can be measured via FRET.

### Establishment of the SMS assay

Cer possesses pro-apoptotic properties [66, 67]. Consistently, we have previously demonstrated that exposure of human cells to high concentrations (10 µM) of α-NH_2_-ω-N_3_-C_6_-Cer for prolonged time periods (24 h) triggers cell death [44]. To identify a Cer concentration not affecting cell viability, we incubated HeLa WT and SMS dK.O. cells [68] with varying concentrations of α-NH_2_-ω-N_3_-C_6_-Cer or the apoptosis inducer camptothecin for 24 h. Next, we determined the total number of adherent cells (**Fig. 2**, a) as well as the proportion of apoptotic cells and/or cells with impaired membrane integrity by annexin V or 7-aminoactinomycin D (7-AAD) staining via flow cytometry (**Fig. S1**, a, b). While concentrations of α-NH_2_-ω-N_3_-C_6_-Cer higher than 5 µM reduced the overall cell number and/or induced apoptosis, we did not detect increased cell death in samples treated with concentrations ≤2.5 µM α-NH_2_-ω-N_3_-C_6_-Cer. Hence, we recommend performing SMS assays in HeLa cells with α-NH_2_-ω-N_3_-C_6_-Cer concentrations ≤2.5 µM.

**Fig. 2.**
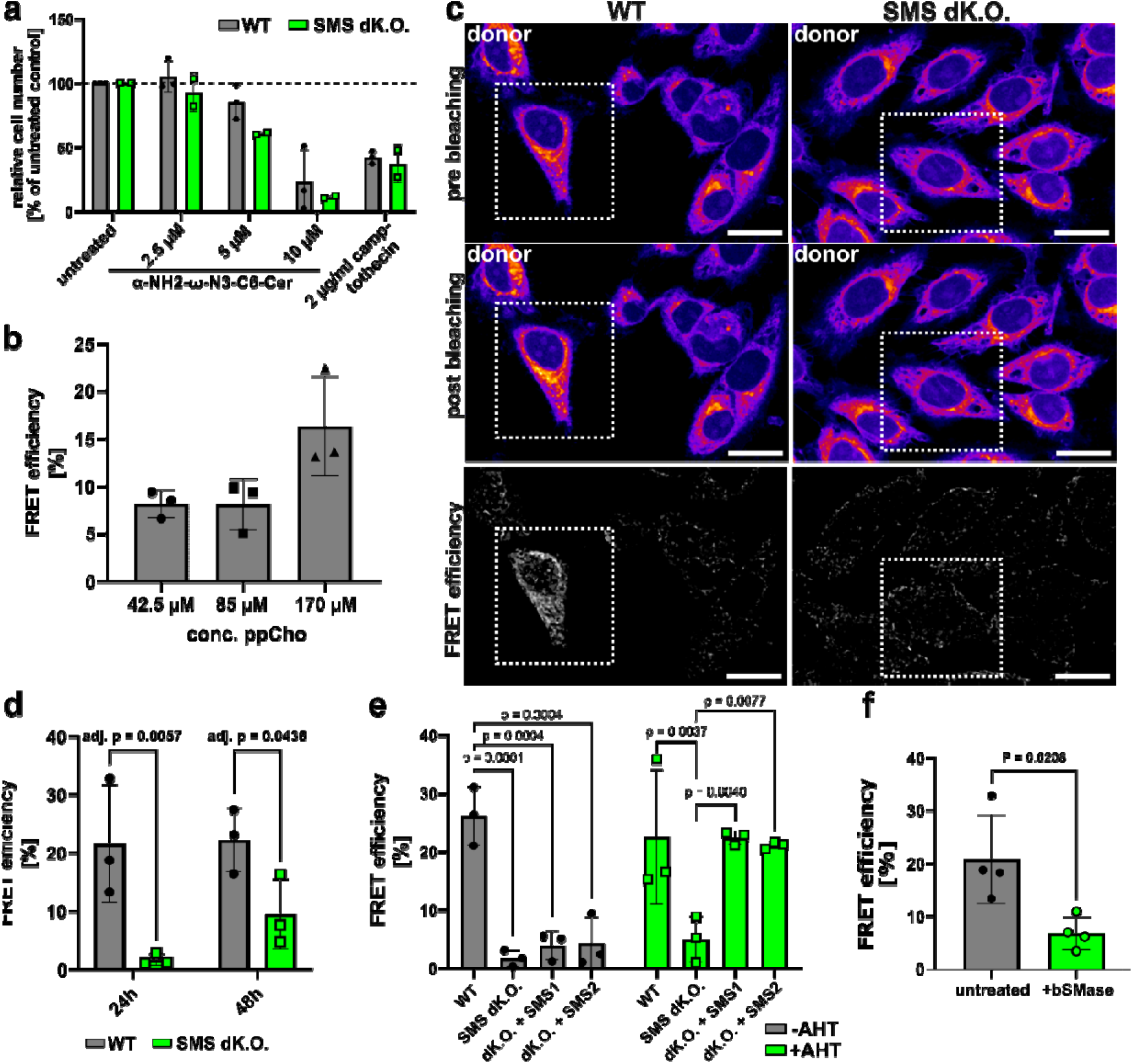
SMS synthase activity can be monitored via FRET. **(a)** Cytotoxic effects of α-NH_2_-ω-N_3_-C_6_-ceramide on HeLa cells. HeLa WT or SMS dK.O. cells were incubated with the indicated concentration of α-NH_2_-ω-N_3_-C_6_-ceramide, camptothecin or left untreated for 24 h. Cells were detached and counted by flow cytometry. Determined cell numbers were normalized to untreated controls (set to 100%). n= 3 (WT) or n=2 (SMS dK.O.).**(b)** FRET efficiency depends on ppCho concentration. HeLa WT cells were incubated with 2.5 µM α-NH_2_-ω-N_3_-C_6_-ceramide and the indicated concentrations of ppCho for 24 h. Cells were fixed, permeabilized and stained with BODIPY-FL-DBCO (donor) and AF546-azide (acceptor). FRET efficiency was determined by FRET acceptor bleaching. n= 3. **(c, d)** Absence of SMSs results in decreased FRET efficiency. HeLa WT or SMS dK.O. cells were incubated with 170 µM ppCho and 2.5 µM α-NH_2_-ω-N_3_-C_6_-ceramide for 24 h or 48 h and FRET efficiency was determined by acceptor bleaching. Areas selected for acceptor bleaching are indicated with white rectangles. n=3. Scale bars: 20 µm. **(e)** SM synthesis can be reconstituted by expression of SMS1/2 in SMS dK.O. cells. HeLa WT, HeLa SMS dK.O., HeLa SMS dK.O.+SMS1 and HeLa SMS dK.O.+SMS2 cells were incubated with 170 µM propargyl-choline, 2.5 µM α-NH_2_-ω-N_3_-C_6_-ceramide and, if indicated anhydrous tetracycline (AHT). FRET efficiency was determined by acceptor bleaching. **(f)** bSMase metabolizes *bio*TFSM. HeLa WT cells were incubated with 2.5 µM α-NH_2_-ω-N_3_-C_6_-ceramide and 170 µM ppCho in the presence or absence of 100 ng/mL recombinant bacterial SMase (bSMase) for 24 h and FRET efficiency was determined by FRET acceptor bleaching. n=4. Statistics: Two-way ANOVA and uncorrected Fisher’s LSD (d) or Šídák’s multiple comparison (e), unpaired Student’s t-test (f). All diagrams show mean ± standard deviation.

Next, we tested whether TFSM biosynthesis can be monitored via FRET. We incubated HeLa WT cells with α-NH_2_-ω-N_3_-C_6_-Cer and varying concentrations of ppCho for 24 h. Samples were fixed, and the potentially generated *bio*TFSM was visualized by click dyes. We first stained the samples with BODIPY-FL-DBCO, which reacts with the *bio*TFSM acyl side chain and acts as FRET donor, and then with AZDye546 azide, which reacts with the ppCho head group of *bio*TFSM and serves as FRET acceptor. Subsequently, we determined FRET efficiency by FRET acceptor bleaching (**Fig. 2**, b). The highest FRET efficiency was measured in samples incubated with 170 µM ppCho. By contrast, we did not detect FRET in samples solely incubated with α-NH_2_-ω-N_3_-C_6_-Cer (**Fig. 2**, a), excluding the contribution of unspecifically bound acceptor dye to the overall FRET signal. However, it is noteworthy that not all dyes are suitable for this approach (**Fig. S2, Supp. Note 1**).

To validate that the measured FRET signal originates from *bio*TFSM2 biosynthesis, we either incubated HeLa WT or SMS dK.O. cells with α-NH_2_-ω-N_3_-C_6_-Cer and ppCho for 24 h or 48 h and again performed FRET acceptor bleaching (**Fig. 2**, c, d). FRET efficiencies were significantly lower in SMS dK.O. compared to WT cells, indicating that *bio*TFSM2 generation is impaired in the absence of SMS1 and 2. Interestingly, FRET efficiencies increased in SMS dK.O. cells after longer treatment periods.

We next reintroduced SMS1 or SMS2 under an anhydrous tetracycline (AHT)-inducible promotor into the SMS dK.O. background and again measured *bio*TFSM2 biosynthesis after 24 h. Expressing SMS1 or SMS2 restored SM biosynthesis in SMS dK.O. cells, suggesting that both isoenzymes can use the artificial substrates to produce *bio*TFSM2 (**Fig. 2**, e).

We next performed the SMS activity assay with HeLa WT cells in the presence of a bacterial sphingomyelinase (bSMase) and again determined the amount of generated *bio*TFSM2 by FRET acceptor bleaching (**Fig. 2**, f). Treatment with bSMase reduced the detected FRET signal, suggesting that the bSMase degraded synthesized *bio*TFSM2, which further corroborates our methodology.

### Detection of *bio*TFSM2 by lipidomics

To further confirm *bio*TFSM2 formation in our assay, we incubated HeLa WT or SMS dK.O. cells with or without α-NH_2_-ω-N_3_-C_6_-Cer and ppCho for 24 h. The lipids were extracted, and *de novo* synthesized *bio*TFSM2 was unambiguously identified using high-resolution LC-MS/MS with synthetic TFSM2 serving as a reference (**Fig. S3**). In WT cells, we could confirm the generation of *bio*TFSM2 when the two substrates were present (**Fig. 3**, a), while the metabolite was completely absent from SMS dK.O. cells.

**Fig. 3.**
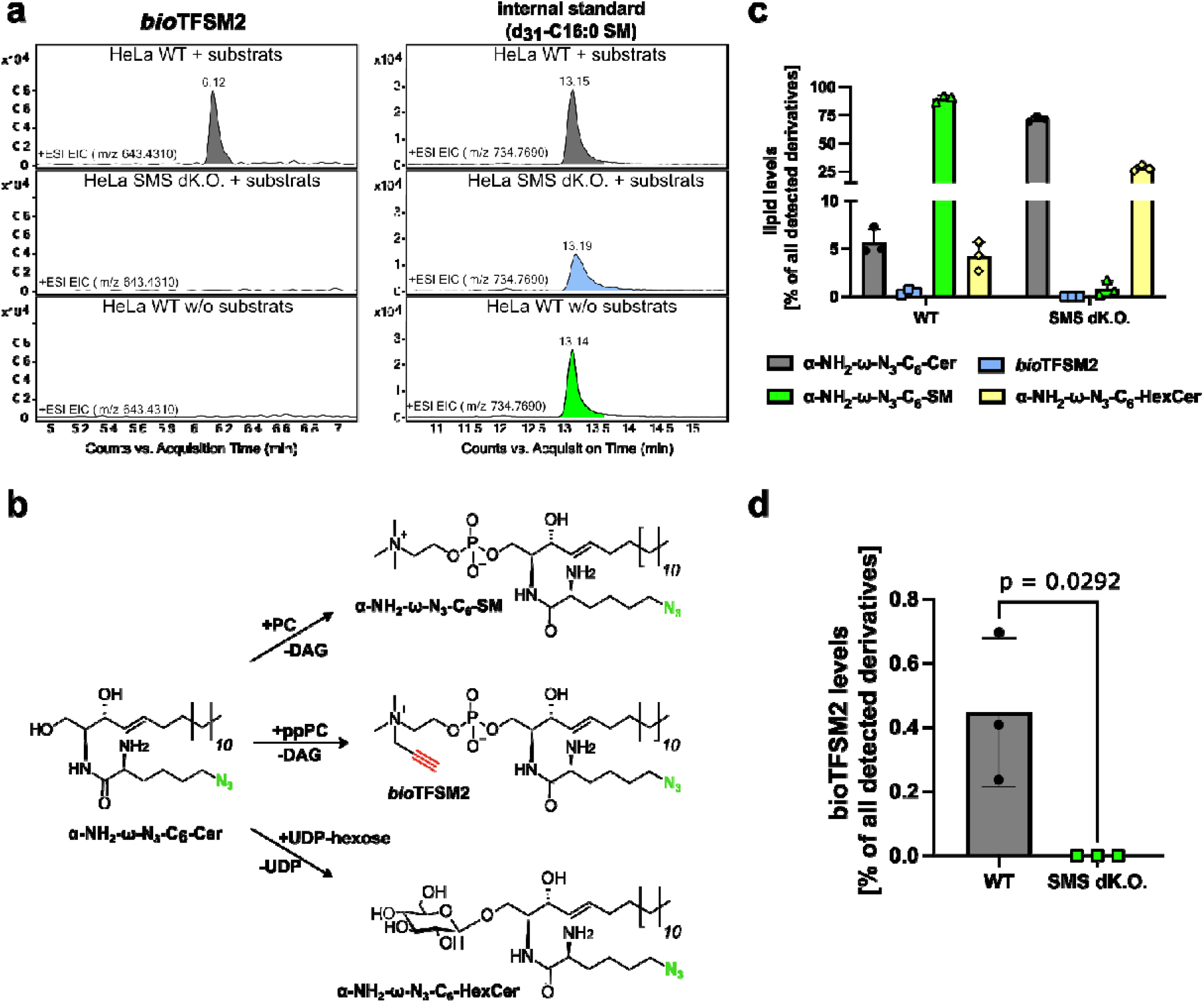
*bio*TFSM2 is synthesized from α-NH_2_-ω-N_3_-C_6_-ceramide and propargyl-choline in living cells. **(a)** *bio*TFSM2 is generated in WT but not SMS dK.O. cells. HeLa WT or SMS dK.O. cells were incubated in the presence (“+ substrates”) or absence (“w/o substrates”) of 2.5 µM α-NH_2_-ω-N_3_-C_6_-Cer and 170 µM ppCho for 24 h. *bio*TFSM2 and the internal standard d_31_-C16:0 SM accounting e.g. for analyte losses during lipid extraction, were detected by LC-MS/MS. +ESI, positive electrospray ionization; EIC, extracted ion chromatogram. **(b)** Biosynthesis of complex sphingolipids derivatives from α-NH_2_-ω-N_3_-C_6_-ceramide. α-NH_2_-ω-N_3_-C_6_-ceramide can be used by SMSs to form *bio*TFSM2 or α-NH_2_-ω-N_3_-C_6_-SM depending on whether ppPC or PC is present. α-NH_2_-ω-N_3_-C_6_-SM possesses a natural phosphocholine head and an α-NH_2_-ω-N_3_-C_6_-ceramide backbone. Alternatively, α-NH_2_-ω-N_3_-C_6_-hexosylceramides (HexCer) can be generated by transferring a hexosyl moiety from UDP-hexoses to α-NH_2_-ω-N_3_-C_6_-ceramide. **(c, d)** Quantification of *bio*TFSM2 metabolites generated during the SMS assay. Quantities of the indicated *bio*TFSM2 metabolites were determined by LC-MS/MS. The proportion of individual metabolites was calculated. n=3. Statistics: unpaired Student’s t-test (d).

In principle, α-NH_2_-ω-N_3_-C_6_-Cer can also be modified with a natural phosphocholine head group as well as with a hexosyl moiety to form hexosylceramide (HexCer), which we refer to as α-NH_2_-ω-N_3_-C_6_-SM or α-NH_2_-ω-N_3_-C_6_-HexCer, respectively (**Fig. 3**, b). Hence, we quantified the formation of these metabolites during our assay conditions by LC-MS/MS. In WT cells, α-NH_2_-ω-N_3_-C_6_-Cer was predominantly metabolized to α-NH_2_-ω-N_3_-C_6_-SM (**Fig. 3**, c), while *bio*TFSM2 only accounts for ~0.5% of all detected artificial lipid derivatives (**Fig. 3**, d). In SMS dK.O. cells, *bio*TFSM2 was completely absent and α-NH_2_-ω-N_3_-C_6_-SM levels were strikingly lower when compared to WT.

Consistent with previous observations [63], generation of α-NH_2_-ω-N_3_-C_6_-HexCer was enhanced in SMS dK.O. cells.

Taken together, we detected the formation of *bio*TFSM2 exclusively in the presence of cellular SMSs, which correlates with our FRET measurements (**Fig. 2**, d), even though α-NH_2_-ω-N_3_-C_6_-SM is predominantly generated from α-NH_2_-ω-N_3_-C_6_-Cer.

#### *Chlamydia trachomatis* synthesizes *bio*TFSM2

The obligate intracellular pathogen *C. trachomatis* depends heavily on host-derived sphingolipids, which are acquired and subsequently incorporated into bacterial membranes [42–47]. We previously demonstrated that exogenously added synthetic TFSMs are metabolized in *C. trachomatis* inclusions [45, 60, 61], whereas other studies have suggested that the pathogen expresses genes for SM biosynthesis [48, 49]. To reconcile these seemingly conflicting findings, we aimed to dissect SM catabolic and anabolic processes during *C. trachomatis* infection.

To monitor SM degradation, we infected HeLa cells in the presence of synthetic TFSM2, determined FRET efficiencies (**Fig. 4,** a) in inclusions (white circle) or a host cell area (blue rectangle) at 32 h post infection (p.i.) and calculated the FRET efficiency ratio “inclusion vs. host cell”. FRET efficiencies in inclusions were strongly reduced compared to host membranes indicating that *C. trachomatis* induces TFSM2 metabolization (**Fig. 4**, b), consistent with our previous observations [45, 60, 61].

**Fig. 4.**
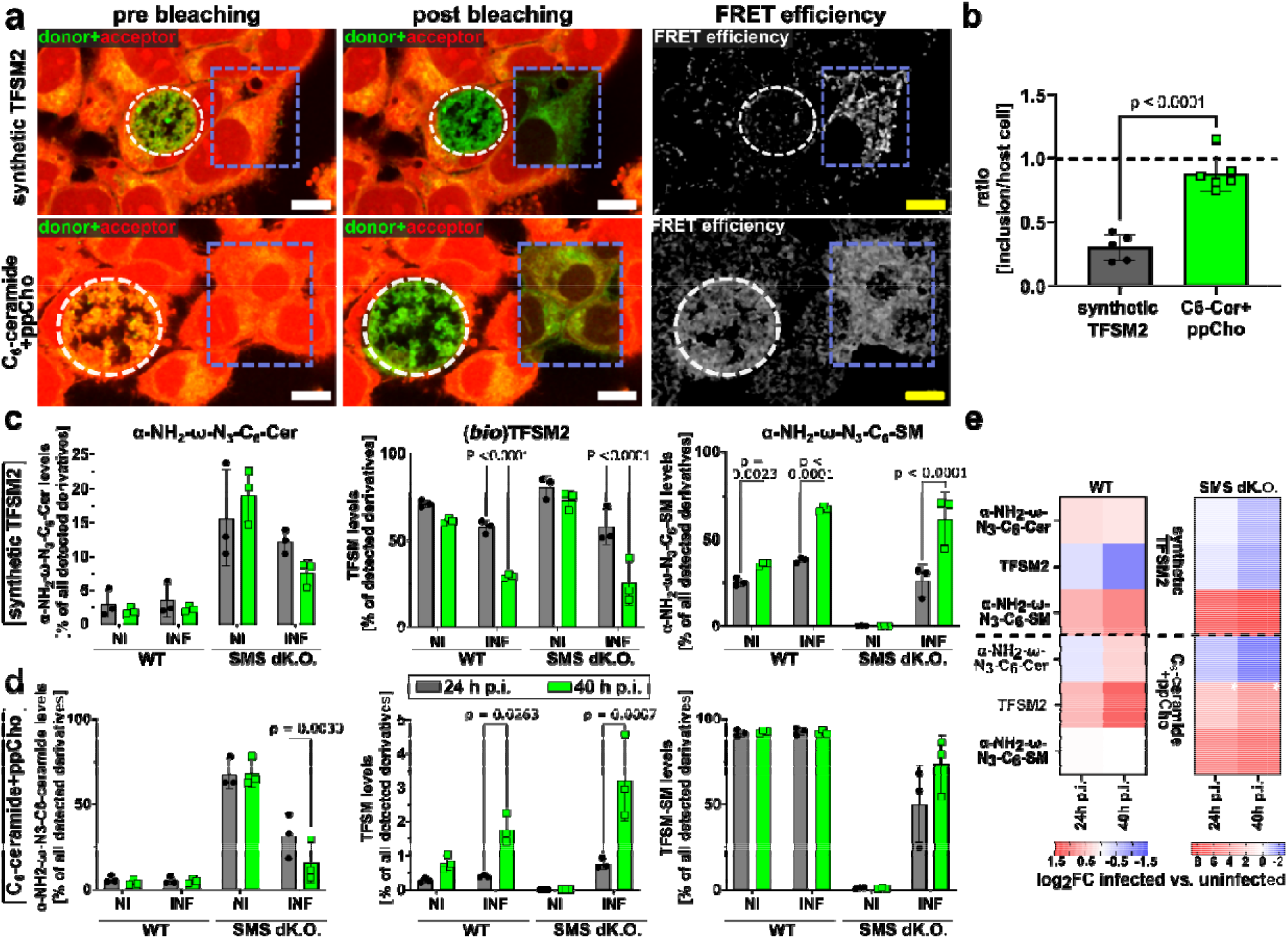
Monitoring TFSM2 degradation and *de novo* synthesis during *C. trachomatis* infection. **(a, b)** Microscopic determination of (*bio*)TFSM2 turnover and biosynthesis. HeLa cells were infected with *C. trachomatis* in the presence of synthetic TFSM2 or α-NH_2_-ω-N_3_-C_6_-ceramide+ppCho for 32 h. FRET efficiencies in inclusions and host cell areas were determined by FRET acceptor bleaching. Scale bars: 5 µm (a). The graphs represent mean values of calculated FRET efficiency ratios “inclusions vs. host cells”. n ≥5. (b). **(c-e)** Analyzing the metabolization of functionalized lipid derivatives during *C. trachomatis* infection by LC-MS/MS. HeLa WT or SMS dK.O. cells were infected with *C. trachomatis* in the presence of synthetic TFSM2 (c) or α-NH_2_-ω-N_3_-C_6_-ceramide+ppCho (d). After the indicated time, lipids were extracted and functionalized lipid derivatives [α-NH_2_-ω-N_3_-C_6_-ceramide, (*bio*)TFSM2, α-NH_2_-ω-N_3_-C_6_-SM, α-NH_2_-ω-N_3_-C_6_-HexCer (not shown)] were quantified by LC-MS/MS. Levels of individual TFSM2 derivatives are shown proportionally to the sum of all detected functionalized lipid derivatives. Log_2_foldchanges (log_2_FC) of infected (INF) vs. uninfected (NI) samples were determined (e). Fields marked with a white asterisk contained zero values in uninfected samples. Statistics: unpaired Student’s t-test (b), Two-way ANOVA, and Šídák’s multiple comparison (c, d).

Next, we repeated the infection experiment in the presence of α-NH_2_-ω-N_3_-C_6_-Cer + ppCho and measured *bio*TFSM2 synthesis by FRET acceptor bleaching. “Inclusion vs. host cell” ratios were significantly higher than those measured with of synthetic TFSM2, suggesting a complex SM metabolization within inclusions.

To further characterize SM metabolism during *C. trachomatis* infection, HeLa WT or SMS dK.O. cells were infected in the presence of synthetic TFSM2 or α-NH_2_-ω-N_3_-C_6_-Cer + ppCho for 24 h or 40 h. Functionalized lipid metabolites were quantified by LC-MS/MS (**Fig. 4**, c, d) and log_2_foldchanges (log_2_FCs) between infected and uninfected samples were calculated (**Fig. 4**, e). Exogenously added synthetic TFSM2 was extensively metabolized during infection, accompanied by α-NH_2_-ω-N_3_-C_6_-SM generation (**Fig. 4**, c and e). These findings suggest that *C. trachomatis* metabolizes TFSM2 to α-NH_2_-ω-N_3_-C_6_-Cer, which is subsequently resynthesized to α-NH_2_-ω-N_3_-C_6_-SM, consistent with our previous observations using a similar sphingolipid derivative [61]. Strikingly, this TFSM2-to-α-NH_2_-ω-N_3_-C_6_-SM conversion was also measured in infected SMS dK.O. cells, whereas α-NH_2_-ω-N_3_-C_6_-SM was completely absent from uninfected cells.

In HeLa WT cells incubated with α-NH_2_-ω-N_3_-C_6_-Cer + ppCho, infection led to increased *bio*TFSM2 levels without affecting α-NH_2_-ω-N_3_-C_6_-SM (**Fig. 4**, d and e). In infected SMS dK.O. cells, we measured enhanced *bio*TFSM2 and α-NH_2_-ω-N_3_-C_6_-SM levels, whereas α-NH_2_-ω-N_3_-C_6_-Cer was markedly decreased when compared to uninfected controls. Together, these results demonstrate that *C. trachomatis* expresses enzymes capable of SM synthesis, in line with previous reports [48, 49].

### Monitoring sphingomyelin metabolism in individual bacteria within *C. trachomatis* inclusions via ExM

Resolving individual chlamydial particles within inclusions by conventional confocal microscopy is challenging, due to the high bacterial density and small size of EBs (Ø ~200-300 nm). The primary amino group present in α-NH_2_-ω-N_3_-C_6_-Cer, (*bio)*TFSM2 and their metabolites enables efficient fixation and makes these probes compatible with ExM [44, 45, 60]. Improved spatial resolution was demonstrated by comparing mitochondria in samples recorded with conventional confocal microscopy (**Fig. S4**, a and b) with those analyzed by 4xExM (**Fig. S4**, c and d). While mitochondria can be localized within unexpanded samples, 4xExM even enabled visualization of cristae in individual mitochondria. Thus, ExM is an ideal tool to study *C. trachomatis* infections.

To quantify *bio*TFSM2 levels at single bacterium level, we infected cells in the presence of α-NH_2_-ω-N_3_-C_6_-Cer + ppCho and analyzed the sample by 4xExM (**Fig. 5**, a-c). During imaging, enlarged particles were observed in some inclusions (**Fig. S5**). We hypothesize that α-NH_2_-ω-N_3_-C_6_-Cer might induce the formation of aberrant bodies, a chlamydial persistent form [69]. Inclusions that were severely affected (**Fig. S5**, a and b) and the enlarged particles themselves were excluded from further analysis.

**Fig. 5.**
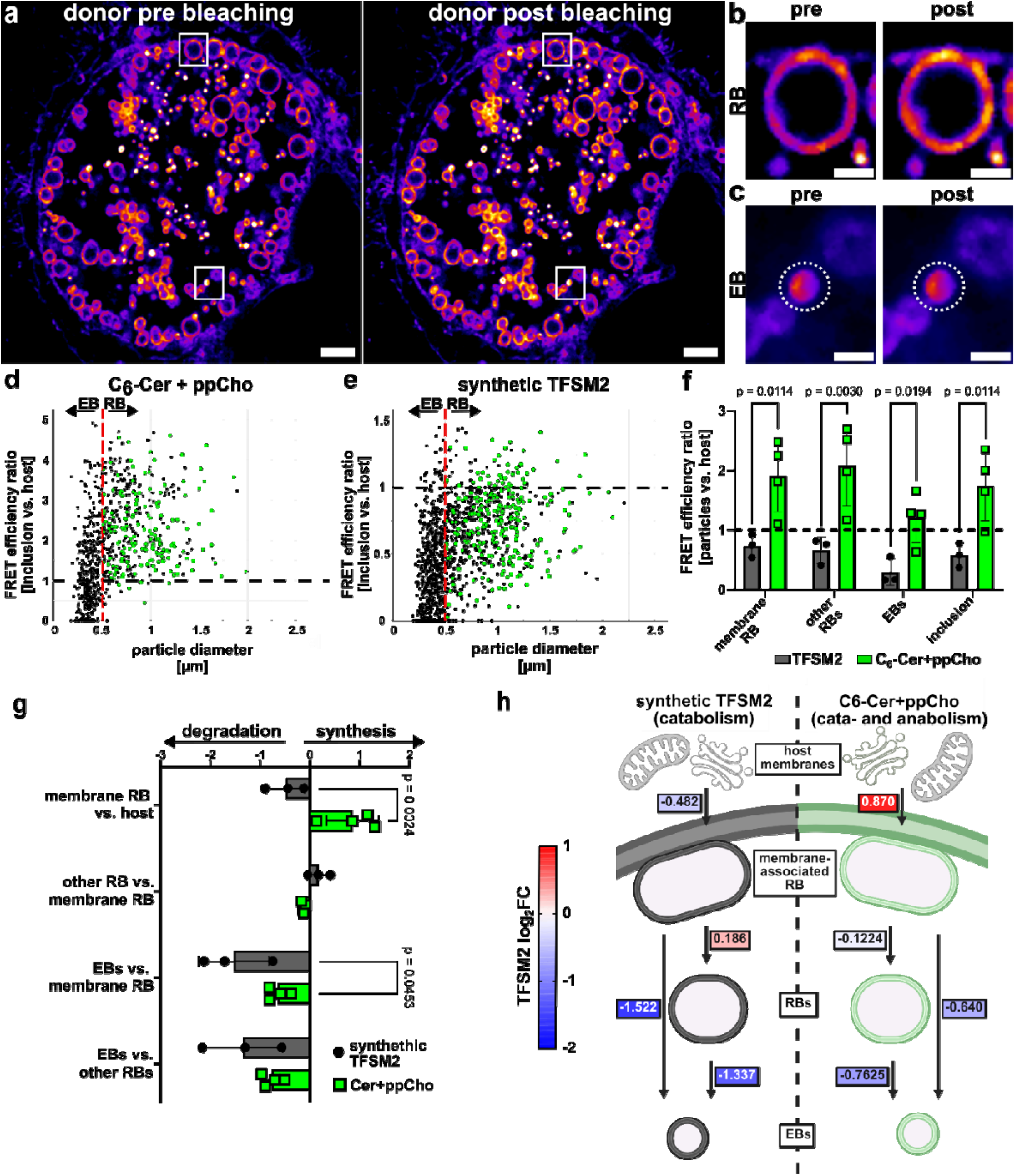
Dissecting TFSM metabolization within *C. trachomatis* inclusions. **(a-f**) Measuring FRET efficiency in individual bacterial particles. HeLa cells were infected with *C. trachomatis* in the presence of α-NH_2_-ω-N_3_-C_6_-ceramide+ppCho (a-d) or TFSM2 (e) for 32 h, 4-fold expanded, and the FRET efficiency of individual bacteria was determined by acceptor bleaching. Particles were categorized as EBs or RBs based on their size (threshold 0.5 µm; d, e) and their association with the inclusion membrane. The average FRET efficiency of each particle class was determined and normalized to the FRET efficiency detected in the corresponding host cell area (f). **(g, h)** Determining TFSM2 metabolization within *C. trachomatis* inclusions. Ratio of average FRET efficiencies of indicated particle classes was calculated and log_2_ transformed. Negative log2FCs indicate a decrease in TFSM2 content (= TFSM2 degradation), while positive log2FCs indicate an increase in the TFSM2 content (= TFSM biosynthesis) between the indicated particle classes. n≥4. Average log2FCs were used to generate a color-coded overview of TFSM2 biosynthesis and metabolization during *C. trachomatis* infection (h). Statistics: Two-way ANOVA and Holm-Šídák’s multiple comparison (f, g).

To quantify *bio*TFSM2 content within individual bacteria, we conducted acceptor bleaching of 4-fold expanded samples (**Fig. 5**, a). In general, RBs (**Fig. 5**, b) showed a higher FRET efficiency than EBs (**Fig. 5**, c), indicated by a lower increase in donor intensity upon bleaching.

FRET efficiencies measured in individual particles were normalized to those of host cell membranes (set to 1). Since RBs (~1 µm) are larger than EBs (<500 nm), these developmental forms could be distinguished by their size. Therefore, normalized FRET efficiencies were plotted against the particle diameter (**Fig. 5, d)**. RBs were further classified as either attached to the inclusion membrane (“membrane RBs”) or located within the inclusion lumen (“other RBs”). As described previously [61], parallel experiments were performed with cells incubated with synthetic TFSM2 to assess TFSM2 degradation within *C. trachomatis* inclusions (**Fig. 5**, e). Mean FRET efficiencies were then calculated for each particle class under both conditions (**Fig. 5**, f).

In samples monitored for *bio*TFSM2 *de novo* synthesis, FRET efficiencies detected in RBs were even higher than in host cell areas, indicating that these particles actively synthesize *bio*TFSM2. In contrast, RBs possessed slightly decreased TFSM2 levels compared to host cell membranes in samples incubated with synthetic TFSM2, indicating that RBs predominantly take up native TFSM2 from the host. In both conditions, EBs exhibited lower FRET efficiencies than RBs, suggesting that TFSM2 is metabolized during RB-EB redifferentiation.

To analyze the dynamics of (*bio*)TFSM2 metabolization, we calculated FRET efficiency log_2_FCs of individual particle classes or host cell membranes (**Fig. 5**, g and h). In both conditions, *(bio*)TFSM2 was not or only marginally altered in RBs detached from the inclusion membrane when compared to membrane RBs. In both experimental settings, EBs contained decreased levels of (*bio*)TFSM when compared to membrane RBs. However, this decrease was more pronounced in samples incubated with synthetic TFSM2 (log_2_FC = −1.522) compared to samples treated with α-NH_2_-ω-N_3_-C_6_-Cer + ppCho (log_2_FC = −0.64).

### *bio*TFSM2 de novo synthesis can be monitored with subcellular resolution in uninfected cells

Next, we performed FRET acceptor bleaching of uninfected host cells in 4-fold expanded samples incubated with α-NH_2_-ω-N_3_-C_6_-ceramide and ppCho. We compared donor fluorescence intensities in whole cells and individual compartments such as the Golgi, mitochondria, the plasma or the nuclear membrane before and after acceptor bleaching (**Fig. 6**, a-e). While the donor signal significantly increased in the Golgi (**Fig. 6**, b) and the plasma membrane (**Fig. 6**, d) upon acceptor bleaching, there was only a small to moderate increase detected in mitochondria (**Fig. 6**, c) and the nuclear envelope (**Fig. 6**, e).

**Fig. 6.**
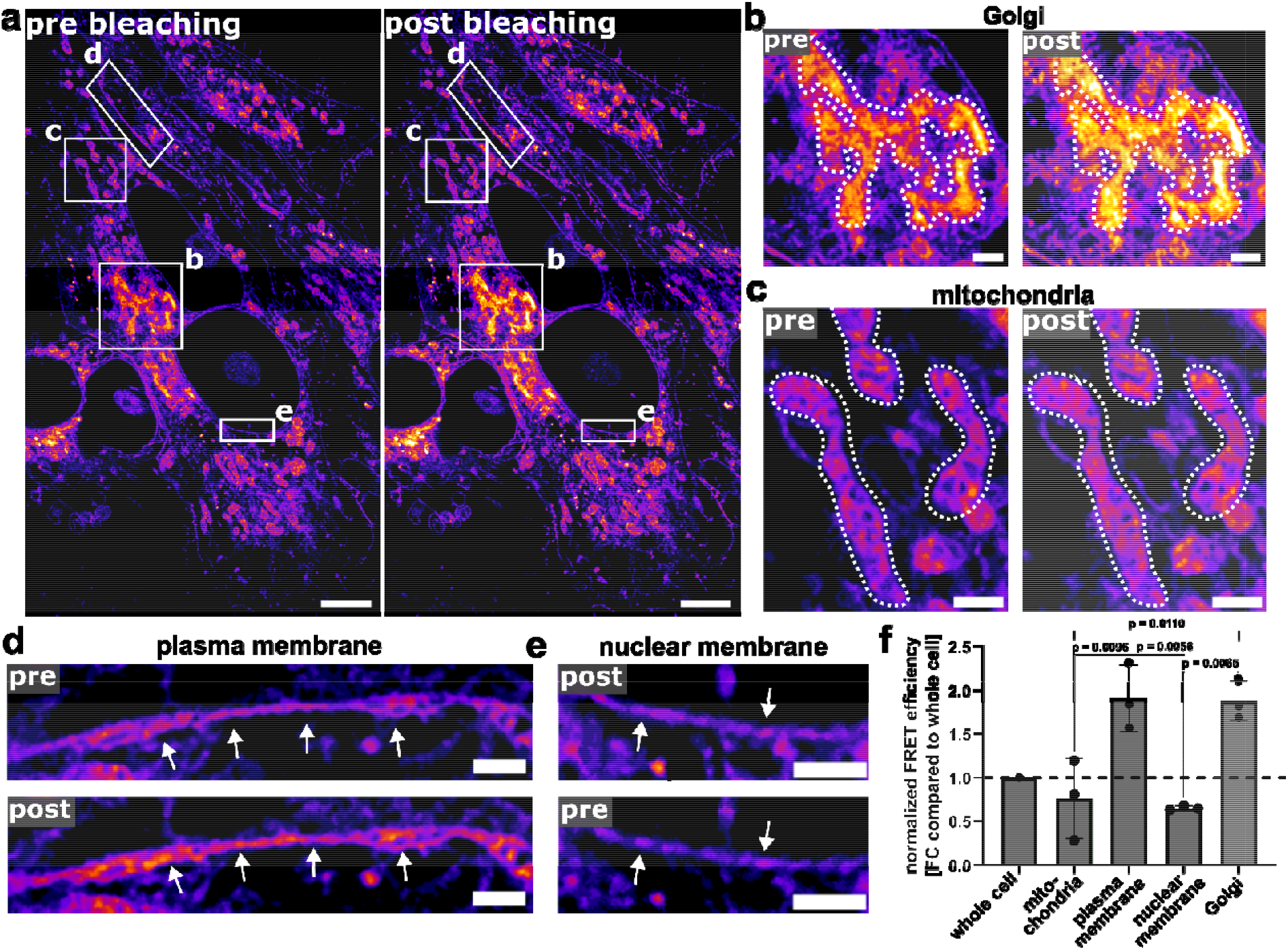
Measuring subcellular distribution of *bio*TFSM via 4xExM. **(a-e)** HeLa cells were incubated with α-NH_2_-ω-N_3_-C_6_-Cer and ppCho and visualized by 4xExM. Donor intensity before (pre) and after (post) bleaching the acceptor fluorophore was compared in whole cells (a) or indicated cellular compartments (b-e). **(f)** FRET efficiency in cellular compartments was determined and normalized to the FRET efficiency detected in the whole cell (set to 1, f). n=3. Scale bar (corrected for 4-fold expansion factor): 5 µm (a) and 1 µm (b-e). Statistics: One-way ANOVA and Tukey’s multiple comparison (f).

To quantify the distribution of *bio*TFSM2, we determined the FRET efficiency in the cellular compartments and normalized it to the FRET efficiency measured in the whole cell (**Fig. 6**, f). As expected, we detected the highest FRET efficiency in Golgi and plasma membrane, the sites of SM biosynthesis [16–18, 64], while mitochondria and the nuclear envelope possess lower FRET efficiencies, suggesting a lower *bio*TFSM2 content. These results demonstrate that our approach enables compartment-specific analysis of SM synthesis.

## DISCUSSION

We present a novel approach to determine cellular SMS activity with spatial resolution. Using artificial substrates, α-NH_2_-ω-N_3_-C_6_-Cer and ppCho, *bio*TFSM2 can be generated in living cells.

The FRET-based readout benefits from the design of (*bio*)TFSM2, as it can be fixed by glutaraldehyde due to its primary amino function, whereas most lipids are lost during permeabilization [44, 45, 65]. This selective retention removes other ppCho-containing lipid species that could otherwise interfere with FRET measurements. Consistently, ppCho-containing lipids are not retained in samples during another lipid ExM protocol [59], unless fixed with a specialized molecular anchoring reagent. This was also demonstrated by the negligible FRET efficiencies measured in SMS dK.O. cells after incubation for 24 h with the substrates (**Fig. 2**, d). The increased FRET efficiency at later times points in SMS dK.O. cells may result from α-NH_2_-ω-N_3_-C_6_-Cer metabolization, leading to the release and incorporation of α-NH_2_-ω-N_3_-hexanoic acid into other lipid species. These modified lipids are retained during permeabilization and thus contribute to the measured FRET signal.

Interestingly, we detected ~20% FRET efficiency in HeLa WT cells, despite a very low *bio*TFSM2 vs. α-NH_2_-ω-N_3_-C_6_-SM ratio, measured by mass spectrometry (**Fig. 3**, c). α-NH_2_-ω-N_3_-C_6_-SM can be fixed and stained with the donor dye, but in contrast to *bio*TFSM2, it cannot be labelled with an acceptor fluorophore. The resulting excess of “silent” donor fluorophores of α-NH_2_-ω-N_3_-C_6_-SM would be expected to dilute FRET signals originating from the few *bio*TFSM2 molecules. However, we detected strongly reduced FRET efficiencies in HeLa SMS dK.O. cells (**Fig. 2**, d and e) and HeLa WT cells treated with bSMase, indicating that the observed FRET signal predominantly originates from *bio*TFSM2. Lipid molecules are densely packed within cellular membranes and SM has been proposed to clusters within lipid microdomains [5–9], which could result in high local *bio*TFSM2/α-NH_2_-ω-N_3_-C_6_-SM concentrations. Their close proximity may promote intermolecular FRET between α-NH_2_-ω-N_3_-C_6_-SM donors and *bio*TFSM2 acceptors. Moreover, we used a low concentration of the donor click dye and thus, only a certain percentage of *bio*TFSM2/α-NH_2_-ω-N_3_-C_6_-SM molecules contains a donor fluorophore, while most acceptor sites are likely occupied. The effective donor population is further limited by donor excitation conditions, as only a subset of donor molecules is excited at low laser intensities. Additionally, dense packaging of donor fluorophores within a membrane can result in self-quenching, which would compensate for the excess of donor dye. These factors likely amplify the FRET signal originating from *bio*TFSM2. Similar effects have been observed previously with lipid FRET probes by us [45] and others [70, 71].

Previous approaches utilized Cer analogs with bulky pre-attached fluorophores [50–53] and it remains unclear how the large residues influence the enzymatic turnover of these analogues. In contrast, our strategy uses minimally modified molecules that allow attachment of the larger fluorophores via click chemistry after the enzymatic reaction has occurred. This design preserves substrate affinity for cellular enzymes, demonstrated by the almost complete conversion of α-NH_2_-ω-N_3_-C_6_-Cer to *bio*TFSM2 or α-NH_2_-ω-N_3_-C_6_-SM in our assay (**Fig. 3**, c). Consistent with this, we previously showed that the staphylococcal bSMase β-toxin can metabolize TFSM, while another SM-FRET probe with pre-attached fluorophores was not recognized as substrate [45]. High substrate affinity is especially important when the inhibitory capacity of competitive inhibitors is determined.

In contrast to other SMS activity assays that rely solely on mass spectrometry [53], the approach we present can, in principle, be performed using standard confocal microscopy. Moreover, the assay enables monitoring *bio*TFSM2 generation with spatial resolution, for instance when determining subcellular distribution of *bio*TFSM2 (**Fig. 6**). We measured significantly higher FRET efficiencies in the Golgi apparatus and the plasma membrane, sites known for containing SMSs [16–18], than in other cellular compartments, further validating the functionality of our assay. Previous studies visualized SM within the Golgi or in the plasma membrane by SM-binding proteins [72, 73]. However, demonstrating biosynthesis of SM in the Golgi required isolation of the respective membranes and detection of metabolized lipid analogs via mass spectrometry [74]. Our method therefore provides a more accessible and spatially resolved alternative.

Unlike other super-resolution imaging techniques, such as Structured Illumination Microscopy (SIM, [75]) or Stimulated Emission Depletion (STED, [76]), which require specialized equipment, 4xExM can yield a spatial resolution of ~60 nm using a common confocal microscope. Other highly specialized imaging techniques, such as matrix-assisted laser desorption/ionization mass spectrometry imaging (MALDI-MSI), enable visualization of natural sphingolipid metabolites, but are limited to single-cell resolution [77].

However, quantitative FRET measurements in 4xExM remain challenging. First, the fluorescence signals are strongly diluted during the expansion process (by factor of 4 for each dimension, 4^3^=64-fold), rendering samples highly susceptible to photo bleaching. This is especially problematic during acceptor bleaching, as the donor fluorescence may be affected during recording the donor channel before the bleaching, leading to an underestimated FRET efficiency. Secondly, the click dyes that we use for staining fixed lipid analogs can introduce unspecific background staining. This is especially challenging during ExM, as higher dye concentrations are required to yield a sufficient fluorescent signal. Hence, quantification of FRET by 4xExM requires a high sample quality, which mainly depends on sufficient incorporation of the lipid analogs in cellular membranes, retention upon permeabilization, efficient staining and embedding into the hydrogel. For detecting (*bio*)TFSM, we recommend using BODIPY-FL-DBCO and AZDye546 azide as FRET pair (see **Supp. Note 1** and **Fig. S2**).

By combining the assay’s spatial resolution with mass spectrometry detection of TFSM2 metabolites, we elucidated the complex SM metabolism in *C. trachomatis* inclusions. When using α-NH_2_-ω-N_3_-C_6_-Cer and ppCho, we measured enhanced FRET efficiency in RBs, that are associated with the inclusion membrane (**Fig. 5**, f-h), which demonstrates that *bio*TFSM2 is either synthesis within these particles or acquired from the host cell as suggested previously by recruitment of SMS1 and 2 to the inclusions membrane [46]. However, detection of *de novo* synthesized *bio*TFSM2 in infected SMS dK.O. cells by lipidomics, consistently with previous reports [48, 49], shows that *bio*TFSM2 can be generated by *Chlamydia*. Upon exogenous addition of synthetic TFSM2, RBs also acquired non-metabolized TFSM2 from the host, highlighting the importance of SM for these bacteria.

After RBs detach from the inclusion membrane and differentiate into EBs, (*bio*)TFSM2 is metabolized irrespective of whether it has been synthesized *de novo* or taken up from the host cell, as indicated by the low FRET efficiencies of EBs in both experimental settings. TFSM2 is mainly converted into the α-NH_2_-ω-N_3_-C_6_-SM metabolite as revealed by our lipidomics analysis (**Fig. 4**, b). This process involves initial degradation of TFSM2 to α-NH_2_-ω-N_3_-C_6_-Cer, which is subsequently reutilized for SM synthesis by the transfer of a natural phosphocholine head group. In the presence of ppCho, α-NH_2_-ω-N_3_-C_6_-Cer can alternatively be modified with the artificial ppCho headgroup to form *bio*TFSM2 during this stage of the *Chlamydia* developmental cycle. Consistent with this, FRET measurements in samples treated with α-NH_2_-ω-N_3_-C_6_-Cer+ppCho shows an attenuated (*bio*)TFSM2 degradation during RB-to-EB differentiation when compared to samples treated with synthetic TFSM2 (**Fig. 5**, g, h). This is further supported by the increased *de novo* synthesis of *bio*TFSM2 detected in infected samples by lipidomics.

Taken together, our data indicate that SM biosynthesis plays a dual role during *C. trachomatis* infection: i.) within RBs associated with the inclusion membrane, that might get fueled by recruitment of the ceramide transfer protein (CERT) to the inclusion [46, 78] and, ii.) during RB-to-EB redifferentiation, which is associated with TFSM2-to-α-NH_2_-ω-N_3_-C_6_-Cer-SM conversion. In line with our previous observations [61], our data provided multiple lines of evidence that SM is acquired from the host, is transiently metabolized to Cer and subsequently is resynthesized to SM, which is the sphingolipid species accumulating in EBs. SM biosynthesis must be mediated by chlamydial factors as formation of α-NH_2_-ω-N_3_-C_6_-SM and *bio*TFSM2 was not affected by the absence of the host SM biosynthesis machinery.

We here present a powerful toolbox that combines metabolic FRET tracking, super-resolution ExM and lipidomics with lipid analogs for monitoring SM biosynthesis and degradation. This approach offers novel possibilities for studying SM metabolism in unprecedented spatial and molecular resolution.

## MATERIALS AND METHODS

### Functionalized lipids

α-NH_2_-ω-N_3_-C_6_-ceramide [44] and trifunctional sphingomyelin 2 (TFSM2, [45]) were synthesized by the group of Jürgen Seibel, Julius-Maximilians-University Würzburg, Germany.

### Cell culture

HeLa wild-type (WT; ATCC^®^ CCL-2™) and SMS-KO cells [63] and HeLa 229 cells (ATCC® CCL-2.1™), were cultured in RPMI+GlutaMAX™ medium (Thermo Fisher, Cat. No. 72400054) containing 10% (v/v) heat-inactivated (56□°C at 30□min) fetal bovine serum (FBS, Sigma Aldrich. Cat. No. F7524) (*culture medium*). Standard tissue culture procedures, including cultivation in a humidified atmosphere with 5 % CO_2_ (v/v) at 37□°C, were performed to maintain the cells.

### Production of SMS dK.O. cells complemented with SMS1 and SMS2

Complementation of sphingomyelin synthase 1 (SMS1) was performed in SMS dK.O. HeLa cells as previously described with modifications [63, 68]. HeLa SMS dK.O. cells were stably transduced with a tetracycline-inducible lentiviral construct pInducer20-FLAG-SMS1. The plasmid and the cell lines (SMS dK.O. and SMS2 complemented cell line) were a kind gift from Joost Holthuis [63, 68].

For lentivirus production, 5 × 10□ low-passage HEK293 cells were seeded per well in a 6-well plate in DMEM supplemented with 10% (v/v) heat-inactivated FBS. Prior to transfection, 12.5 µM chloroquine (Sigma-Aldrich) was added to the culture medium. Lentiviral plasmid mixtures containing 4 µg total DNA (pInducer20-FLAG-SMS1) together with the packaging plasmids psPAX and pVSV-G were prepared in 500 mM CaCl□. Subsequently, 2× HEPES-buffered saline (50 mM HEPES, 140 mM NaCl, 1.5 mM Na□HPO□, pH 7.05) was added dropwise while bubbling. The transfection mixture was incubated at room temperature for 15 min and then added dropwise to the cells. Culture medium was changed 6 h post-transfection. After 48 h, the lentivirus-containing medium was harvested (0.45 µm filter), mixed 1:1 (v/v) with DMEM containing 8 µg/ml polybrene (Sigma-Aldrich, TR-1003) and used to infect HeLa SMS1/2 DKO cells. At 24 h post-infection, the medium was replaced with DMEM containing 1 mg/mL G418 (Sigma-Aldrich, G8168) to select for stably transduced cells. Selection medium was changed daily for 5-6 days. Following antibiotic selection, stable cell populations were expanded.

### Cultivation of cells to measure the subcellular distribution of *bio*TFSM via 4xExM

HeLa cells were seeded a day prior to the experiment in a 24-well plate (6□× □10^4^ cells per well) on cover slips. The cells were incubated in medium containing 1% FBS, 2.5 µM α-NH_2_-ω-N_3_-C_6_-ceramide and 170 µM propargyl-choline (Jena Bioscience, Cat. No. CLK-066) for 24 h. The cells were washed twice with Dulbecco’s phosphate-buffered saline (DPBS; Gibco, Cat. No. 14190169), and fixed and processed as described below.

### Purification of *C. trachomatis* elementary bodies

For this study, the L2□/434/Bu (ATCC VR-902B) serovar of *C. trachomatis* was used. The purification of *C. trachomatis* was performed as described before [79]. In brief, HeLa 229 cells were infected with *C. trachomatis* at MOI 1, which was then expanded for 48 h. To disrupt and mechanically lyse the cells, glass beads (2.85–3.45□mm, Roth, Cat. No. A557.1) were used. This was followed by centrifugation for 10□min at 755□× □*g* at 4□°C and another centrifugation of the supernatant for 30□min at 40,000□× □*g* at 4□°C to collect the bacteria. To wash the bacteria, 1 x sucrose–phosphate–glutamic acid (SPG) buffer [7.5% sucrose (Roth, Cat. No. 4621.2), 0.052% KH_2_PO_4_ (Roth, Cat. No. 3904.1), 0.122% Na_2_HPO_4_ (Roth, Cat. No. P030.2), 0.072% L-glutamine (Gibco, Cat. No. 25030081)] was used and the centrifugation step was repeated. After resuspending the chlamydial pellet in 1 x SPG buffer, the *Chlamydia* were singularized by passing the suspension through G20 (B. Braun, Cat. No. 612-0141) and G18 (B. Braun, Cat. No. 612-0147) hollow needles. Purified *Chlamydia* were stored at −80 °C. To determine MOI1, titration was performed and PCR was used to confirm that the bacteria were free from Mycoplasma contamination.

### Cultivation of *C. trachomatis* infected cells in the presence of lipid derivatives for microscopy

HeLa cells were seeded a day prior to the experiment on cover slips in a 24-well plate (6□× □10^4^ cells per well). For the infection, the medium was exchanged to fresh *culture medium*, and the cells were infected with *C. trachomatis* at MOI 1. The cells were treated as indicated. TFSM2 (10 µM) and ppCho (170 µM) were added 3 h after infection and medium containing 1% FBS was used, while α-NH_2_-ω-N_3_-C_6_-ceramide (2.5 µM) was added 6 h after the infection. 32 h after infection, the cells were washed twice with DPBS and fixed and processed as described below.

### Cytotoxicity assay

HeLa WT or SMS dK.O. cells were treated with the indicated concentrations of α-NH_2_-ω-N_3_-C_6_-Cer or 2 µg/mL camptothecin (Merck, Cat. No. 208925) for 24 h. Cells were washed thrice with DPBS and detached with trypsin. Subsequently, cells were incubated with 0.85% (v/v) APC annexin V (BD Pharmingen, Cat. No. 550475), 0.85% (v/v) 7-AAD (BD Pharmingen, Cat. No. 559925), 2 mM CaCl_2_ (Roth, Cat. No. 5239.1), and 1% FBS in PBS for 10 min/RT. Cells were counted and analyzed for 7-AAD and APC fluorescence at an Attune NxT flow cytometer (Thermo Fisher).

### Sample preparation for fluorescence microscopy

After treatment of cells with lipid analogs, samples were washed twice with DPBS and fixed with 4% PFA in PBS (Morphisto, Cat. No 11762), complemented with 0.25 % glutaraldehyde (Sigma Aldrich, Cat. No. G5882) for 30 min/RT. Samples were washed thrice with DPBS and permeabilized with 0.2% Triton X-100 (Roth, Cat. No. 6909) for 20 min/RT.

For visualization of mitochondria, samples were first incubated with blocking buffer (5% FBS in DPBS) for 1 h and then, with 1:50 anti-Prx3 antibody (OriGene, Cat. No. TA322472) in blocking buffer overnight. Samples were washed thrice with DPBS and incubated with 1:100 anti-rabbit AF994 antibody (ThermoFisher. Cat. No. A-11012) in blocking buffer for 1 h/RT. Samples were fixed with 4% PFA in PBS for 10 min/RT.

Subsequently, samples were washed thrice and incubated with 0.5 µM (conventional microscopy) or 4 µM (4xExM) BODIPY-FL-DBCO (Jena Biosciences, Cat. No. CLK-040-05) in HBSS (Gibco™, Cat. No. 14025-100) for 1 h/37°C at 170 rpm. Samples were washed five times and incubated with 5 µM (conventional) or 20 µM (4xExM) AZDye546-azide (Jena Bioscience, Cat. No. CLK-1283-AZ) or AZDye546-picolyl-azide (Jena Bioscience, Cat. No. CLK-1284-AZ) in click buffer [50 μM CuSO4, 2.5 mM sodium ascorbate (Sigma, Cat. No. A4034), 250 μM Tris(3-hydroxypropyltriazolylmethyl)amine (THPTA; Sigma, Cat. No. 762342) in DPBS] for 1 h/37°C at 170 rpm. Samples were washed thrice with DPBS and then again incubated with DPBS for 3×15 min on a see-saw rocker.

Samples were either mounted with Mowiol [24 g glycerol (Roth, Cat. No. 3783.2), 9.6 g Mowiol^®^ 4-88 (Roth, Cat. No. 0713.2), 48 mL 0.2M TRIS-HCl pH 8.5 (Sigma, Cat. No. T1503), 24 mL Millipore H_2_O] or used for ExM.

### Expansion microscopy

The protocol is based on previous publications [44, 45, 55]. After fluorescence staining, samples were fixed with 0.25% glutaraldehyde in DPBS for 10 min/RT and washed thrice with DPBS. Subsequently, samples were soaked in ~20 µL 4x monomer solution [8.625% sodium acrylate (Sigma, Cat. No. 408220), 2.5% acrylamide (Sigma, Cat. No. A9926), 0.15% N,N’-methylene bisacrylamide (Sigma, Cat. No. 146072), 2 M NaCl (Sigma, Cat. No. S5886) in PBS] on a parafilm. Then, a drop of 56 µL (for Ø12 mm cover slips) 4x monomer solution complemented with 0.2% (w/v) ammonium persulfate (Sigma, A3678) and 2% (v/v) tetramethyl ethylene diamine (TEMED) was pipetted on a parafilm in a petri dish. The cover slip was immediately turned into the monomer solution and incubated in a closed Petri dish for 75 min/RT. Gels were removed from the cover slips and digested with 8 U/mL proteinase K (Sigma, Cat. No. P4860) in digestion buffer [50 mM Tris pH 8.0, 1mM EDTA (Sigma, Cat. No. ED2P), 0.5% Triton X-100 and 0.8 M guanidine HCl (Sigma, Cat. No. 50933)] for 30 min/RT. Gels were incubated in excess of water until the expansion process was completed. Samples were imaged in a poly-L-lysin-coated µ-Slide 1 well glass bottom dish (ibidi, Cat. No. 82107) or on a silanized cover slip in a microscopy cell chamber (ThermoFisher, Cat. No. A7816) that was filled with water.

### FRET efficiency measurements

FRET efficiency in conventional unexpanded samples was measured on a Leica Stellaris 5 confocal microscope using the built-in FRET acceptor bleaching wizard.

For FRET measurements within 4xExM samples, the donor channel (BODIPY-FL) pre-bleaching was recorded. The acceptor (AZDye546) was bleached in the whole region, and the donor channel post-bleaching was recorded. Regions of interest (e.g. individual bacterial particles in an inclusion; ROI) were selected in Fiji [80] and mean fluorescence intensities of the donor channel in individual ROIs were determined before and after bleaching. The FRET efficiency was determined based on the formula:

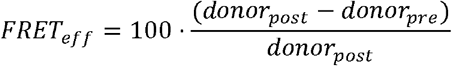

For measurements of individual chlamydial particles, the diameter of bacteria was determined and plotted against the FRET efficiency detected in the particle using R studio[81] and ggplot[82]. Particles were categorized based on their diameter as EBs (<500 nm) or RBs (>500 nm) and their association with the inclusion membrane. Average FRET efficiencies were calculated for individual particle classes and either normalized to the FRET efficiency detected in host cell areas or used to calculate FRET efficiency foldchanges between individual particle classes.

### Silanization of cover slips

First, Ø24 mm cover slips were rinsed with MilliQ water and incubated with MilliQ water and then, with 5 M KOH in an ultrasonic cleaning bath for 15 min each. Cover slips were washed thrice in MilliQ water and further incubated in an ultrasonic cleaning bath with MilliQ water for 15 min and subsequently, with 100% EtOH for 15 min. Cover slips were air dried, covered with ~200 µL silanization solution [90% (v/v) EtOH, 5% (v/v) MilliQ water, 5% (v/v) AcOH, 0.1 % (3-ayminopropyl)triethoxysilane (ThermoFisher, Cat. No. A3648)] and incubated at RT until the liquid was completely evaporated. Cover slips were first dipped in MlliQ H_2_O and then into 100% EtOH and again air dried. This process was repeated 2-3 times until the cover slip was free from residual silane.

### Lipidomics

#### Sample preparation for the detection of bioTFSM2 biosynthesis in HeLa cells

For samples tested for *bio*TFSM2 synthesis (**Fig. 3**, c and d), HeLa WT and SMS dK.O. cells were seeded a day prior to the experiment in a 6-well plate (2.5□×□10^5^ cells/well). The medium of the cells was exchanged to fresh *culture medium*, the cells were treated as indicated and 2.5 µM□µM α-NH_2_-ω-N_3_-C_6_-ceramide and 170 µM propargyl-choline were added respectively. After 24 h, cells were washed 2 × with DPBS, resuspended in 1 mL methanol (LC-MS CHROMASOLV, Fluka analytical, Cat. No. 34966-1L) and subjected to sphingolipid extraction.

For measuring TFSM metabolites during *C. trachomatis* infection (**Fig. 4**, c-e), HeLa WT and SMS-KO cells were seeded a day prior to the experiment in a 6-well plate (24 h time point: 4□×□10^5^ cells per well that will be infected and 2□×□10^5^ cells per well that will not be infected, 40 h time point: 4□×□10^5^ cells per well that will be infected and 1□×□10^5^ cells per well that will not be infected). The next day, the medium of the cells was exchanged to fresh *culture medium*. If indicated, the cells were infected with *C. trachomatis* at MOI 1. 6 h after infection, the medium was exchanged to fresh *culture medium* or medium containing 1% heat-inactivated FBS as indicated. The cells were treated as indicated and 2.5 µM α-NH_2_-ω-N_3_-C_6_-ceramide, 10□µM TFSM2 and 170 µM propargyl-choline were added respectively. At the indicated time points, cells were washed 2-times with DPBS, resuspended in 1 mL methanol and subjected to sphingolipid extraction.

#### Detection of (bio)TFSM2 derivatives by LC-MS/MS

Cell suspensions were first subjected to lipid extraction, which was performed according to a protocol for the determination of canonical cellular sphingolipids [83]. In brief, all samples were extracted with a mixture of methanol/chloroform (2:1, v:v) containing various internal standards (ISTD). The lipid extracts were then chromatographically separated on a 1290 Infinity II HPLC (Agilent Technologies, Waldbronn, Germany), which was equipped with a Poroshell 120 EC-C8 column (3.0 × 150 mm, 2.7 μm; Agilent Technologies). Depending on the objective, detection was carried out either on a 6550 quadrupole time-of-flight mass spectrometer (Agilent Technologies, initial screening for sphingolipid analogues) or a 6495C triple-quadrupole mass spectrometer (Agilent Technologies, targeted quantification). In both cases, precursor ions were generated by positive-mode electrospray ionization (ESI+) and all settings of the MS and MS/MS detector were adopted from a previously described method [83]. The quantification of sphingolipid analogues was performed in multiple reaction monitoring (MRM) mode using the following mass transitions (qualifier transitions in parentheses): *m/z* 436.4 → 264.3 (282.3) for α-NH_2_-ω-N_3_-C_6_-Cer, *m/z* 534.5 → 264.3 (282.3) for C17:0 Cer (ISTD), *m/z* 619.4 → 184.0 (86.1) for α-NH_2_-ω-N_3_-C_6_-SM, *m/z* 643.4 → 208.0 (125.0) for (*bio*)TFSM2, *m/z* 734.6 → 184.0 (86.1) for d_31_-C16:0 SM (ISTD), *m/z* 616.4 → 264.2 (598.4) for α-NH_2_-ω-N_3_-C_6_-HexCer and *m/z* 714.6 → 264.2 (696.6) for C17:0 HexCer (ISTD). Peak areas of sphingolipid derivatives were determined using MassHunter Quantitative Analysis software (version 10.1, Agilent Technologies) and quantification was achieved in relation to their corresponding ISTDs that were concentrated to 0.25 µM in the final extracts.

### Statistical analysis

Statistics were performed in GraphPad Prism. Diagrams represent mean values ± standard deviation.

## ACKNOWLEDGEMENTS

We thank Joost C. M. Holthuis (University of Osnabrück) for providing the HeLa WT and SMS dK.O cells. We thank Markus Sauer for fruitful discussion and Stefan Sachs (JMU Würzburg) for providing protocols for cover slip silanization and general support of expansion microscopy. We would like to acknowledge the assistance of the Core Facility BioSupraMol supported by the DFG. Daniel Herrmann’s technical support during the LC-MS/MS analyses is also greatly appreciated.

## Funding

Deutsche Forschungsgemeinschaft (DFG), Project code 116162193 (Leica TCS SP5 CLSM) and Project code INST93/1159-1 FUGG (Leica Stellaris 5 CLSM).

DFG, RTG2581 (B.K., J.S., T.R., V.K.P.) DFG, SFB 1583 (T.R.).

European Research Council (ERC), Grant No. ERC-2018-ADG/NCI-CAD (T.R.).

Bavarian State Ministry of Science and the Arts and the University of Würzburg to the Graduate School of Life Sciences (GSLS), University of Würzburg funds (M.R.).

## Author contributions

Conceptualization: M.R.

Methodology: M.R., F.W., A.M., F.S., J.F., L.K., and J.S.

Investigation: F.W., A.M., F.S., B.K., T.R., V.K.P., and M.R.

Visualization: M.R.

Supervision: B.K., J.S., T.R., V.K.P., M.R.

Writing—original draft: M.R. Writing—review & editing: all authors

