## Supplemental text and figures for "Monitoring Sphingomyelin Biosynthesis at Nanoscale Resolution by Expansion Microscopy"

### Supplementary Text

Supplementary Note 1

The type of fluorophore used to visualize (*bio*)TFSM can impact the FRET read out. The dyes used for the FRET system should possess the following properties: i.) Penetrate the sample properly thereby enabling high staining efficiency. This is especially important for the DBCO dye as the azide function of the lipid analog is buried in the non-polar membrane bilayer. ii.) The two fluorophores should be able to interact via FRET when they are attached to (*bio*)TFSM. This not only requires overlap of the donor emission and the acceptor excitation spectra but is also affected by the chemical orientation of the two fluorophores to each other [84], which can hardly be predicted. iii.) The background FRET efficiency should be as low as possible. FRET background can be caused by nonspecific binding of the two dyes to the sample. This can either result in the accumulation of the dyes at certain sites in the samples or even in covalent coupling of the two fluorophores via a direct click reaction between the two dye molecules, resulting in "dye dimers". iv.) The dyes should be compatible with ExM.

We recommend using BODIPY-FL DBCO as a donor as BODIPY fluorophores possess non-polar characteristics and were previously used to stain lipids [45, 85].

We tested dye pairs for FRET background (**Supp. Figure 2**, a). Therefore, we incubated HeLa 229 cells infected with *C. trachomatis* either with α-$NH_2$-ω-$N_3$-$C_6$-ceramide and ppCho (*bio*TFSM formation) or solely with α-$NH_2$-ω-$N_3$-$C_6$-ceramide. Then, we either stained with BODIPY-FL DBCO and AZDye546 azide (Cat. No. CLK-1283-AZ) or AF594 DBCO and Atto647N azide. Subsequently, FRET efficiencies in inclusions and host cell areas were determined. If we used BODIPY-DBCO as donor and AZDye546 azide as acceptor, we detected a FRET efficiency of ~15% in samples treated with α-$NH_2$-ω-$N_3$-$C_6$-ceramide and ppCho. The background FRET efficiency measured in samples solely incubated with α-$NH_2$-ω-$N_3$-$C_6$-ceramide was negligible, suggesting that the BODIPY-FL/AZDye546 dye system predominantly detects FRET originating from (*bio*)TFSM. By contrast, we also detected a higher background FRET efficiency in samples incubated with α-$NH_2$-ω-$N_3$-$C_6$-ceramide and stained with AF594/Atto647N. The background was especially pronounced in *C. trachomatis* inclusions.

The AZDye546 azide (Cat. No. CLK-1283-AZ) we used here and previously[45] was discontinued by the manufacturer and replaced by a dye (Cat. No. CLK-1283**A**-AZ) with similar spectral properties but a different chemical structure. We compared the two dyes in combination with BODIPY-FL-DBCO in cells treated with synthetic TFSM2 (**Supp. Figure 2**, b). We detected FRET in the sample stained with the original AZDye546 azide (Cat. No. CLK-1283-AZ), but not with the replacement product (Cat. No. CLK-1283**A**-AZ). However, AZDye546-picolyl-azide (Cat. No. CLK-1284-AZ) possesses a similar chemical structure as the original AZDye546 (Cat. No. CLK-1283-AZ) and only varies in the linker region. We detected a TFSM2 by FRET using the BOIDPY-FL/AZDye546-picolyl dye system. As AZDye546-picolyl-azide is still available, we recommend using this dye instead of the AZDye546 azide replacement product (Cat. No. CLK-1283**A**-AZ). Alternatively, other FRET systems could be used to measure TFSM2 metabolization

or *de novo* synthesis, but it should be carefully tested whether the dye pairs are suitable for this application.
Additionally, BODIPY-FL and AZDye546 are compatible with ExM, since not all fluorophores survive the expansion process [86].

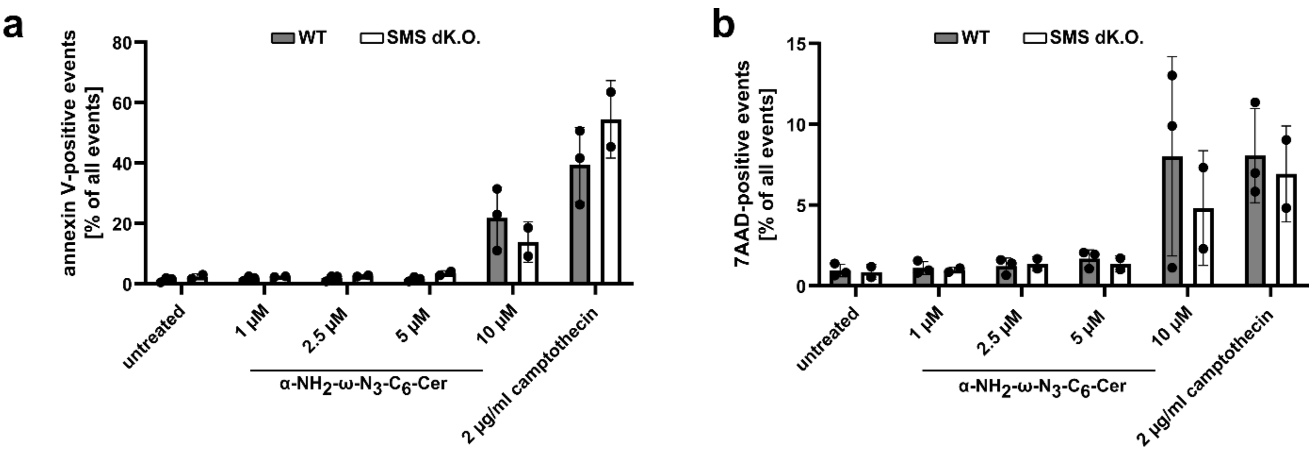


**Fig. S1. High concentrations of α-$NH_2$-ω-$N_3$-$C_6$-Cer induce cell death.** HeLa cells were treated with the indicated concentration of α-$NH_2$-ω-$N_3$-$C_6$-Cer or the apoptosis inducer camptothecin. Cells were stained with fluorescent annexin V **(a)** and 7-AAD **(b)** and analyzed by flow cytometry.

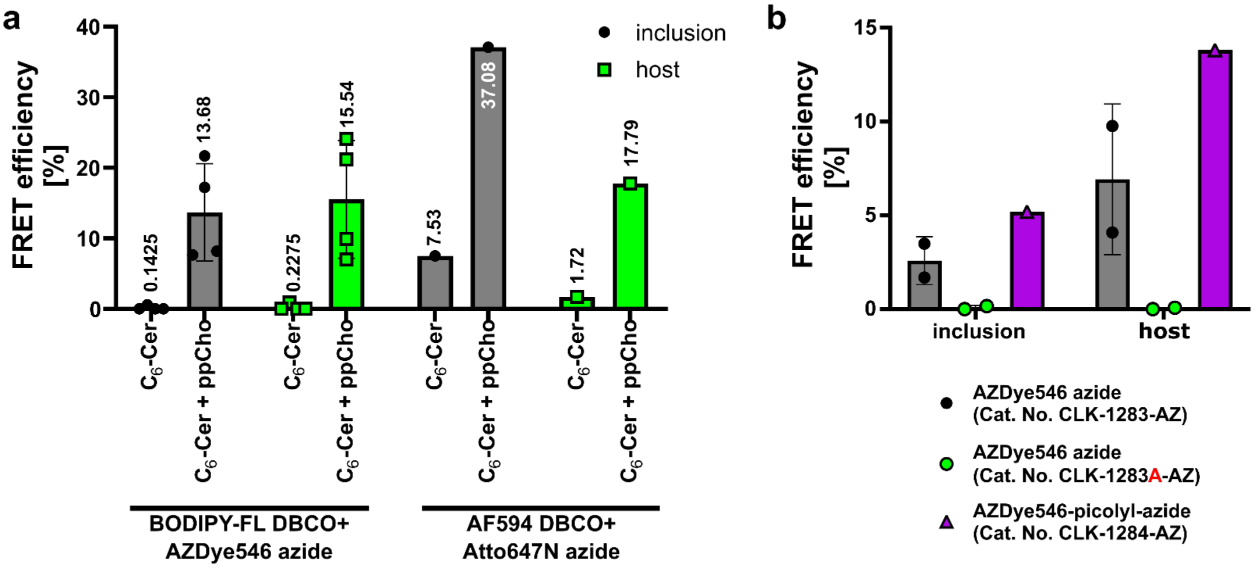


**Fig. S2. Detection of FRET efficiency in sample containing (*bio*)TFSM is affected by the click dyes used.** HeLa cells were infected with C. trachomatis for 32 h and treated with α-$NH_2$-ω-$N_3$-$C_6$-Cer or α-$NH_2$-ω-$N_3$-$C_6$-Cer and ppCho **(a)** as well as TFSM2 **(b)**. Samples were fixed, permeabilized and stained with BODIPY-FL DBCO (a, b) or if indicated with AF594 DBCO. Subsequently, samples were stained with the indicated AZDye 546 azide (a, b) or Atto647N azide (a). FRET efficiencies in inclusions and host cell areas were determined by acceptor bleaching.

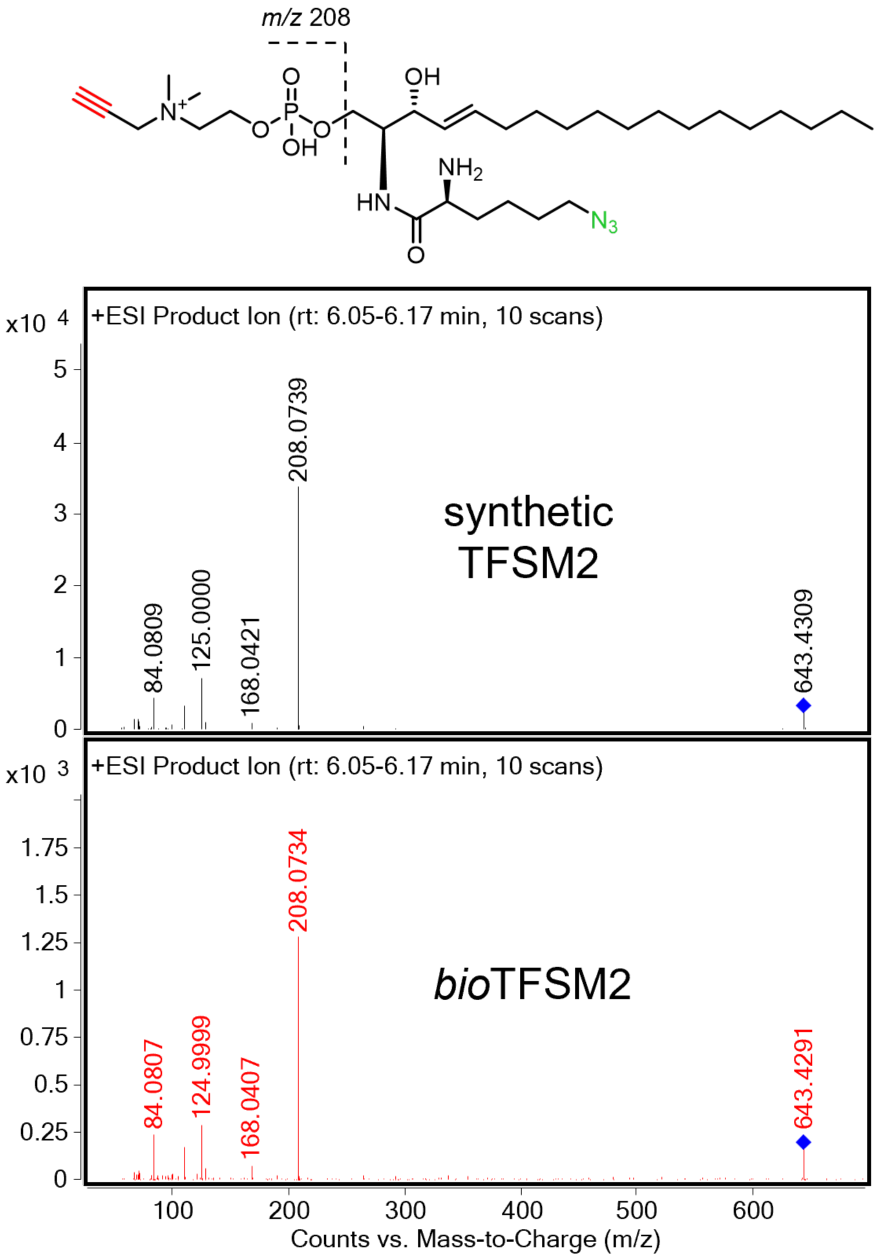


**Fig. S3. Similar MS/MS spectra of synthetic TFSM2 and de novo-formed *bio*TFSM2 were obtained by high-resolution LC-MS/MS.** The fragment ion m/z 208 confirms the presence of a propargyl choline group. +ESI, positive electrospray ionization.

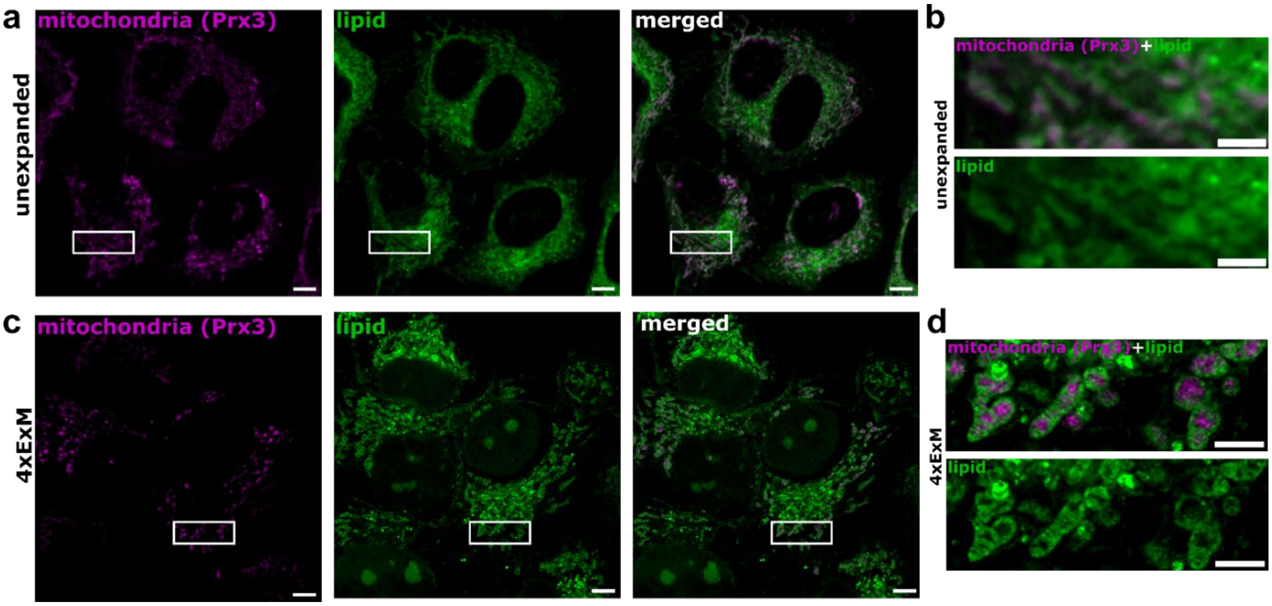


**Fig. S4. Monitoring lipids by ExM.** HeLa 229 cells were incubated with α-$NH_2$-ω-$N_3$-$C_6$-Cer and ppCho, fixed and mitochondria were stained with an Prx3 antibody. Samples were either imaged by conventional confocal laser scanning microscopy **(a, b)** or via 4xExM **(c, d)**. Scale bar: 5 µm (a, c), 2 µm (b, d). Scale bars of expanded samples were corrected for 4x expansion factors.

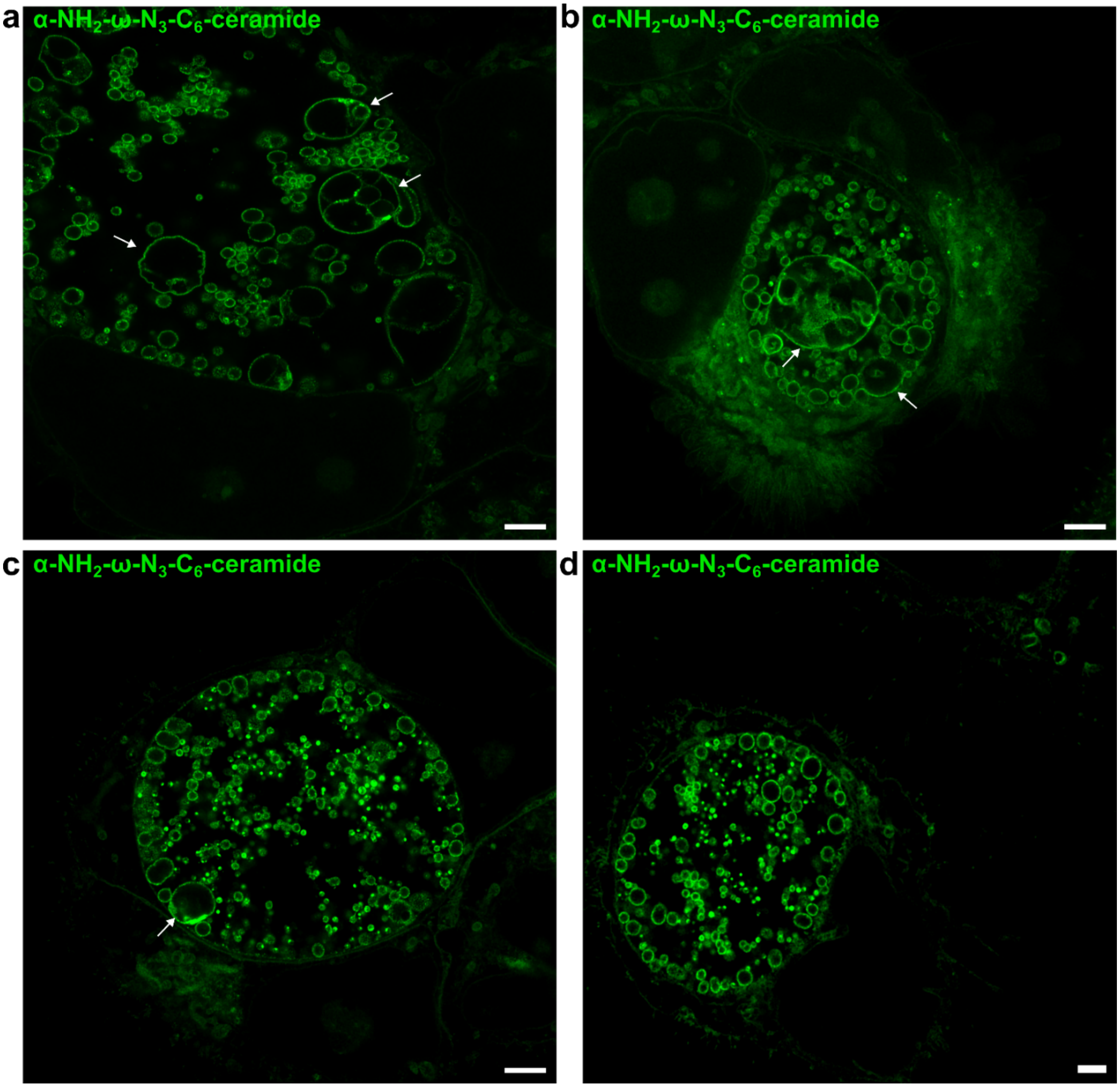


**Fig. S5. α-$NH_2$-ω-$N_3$-$C_6$-ceramide can induce formation of aberrant bodies in a subset of inclusions.** HeLa 229 cells were infected with *C. trachomatis* in presence of α-$NH_2$-ω-$N_3$-$C_6$-ceramide and ppCho for 32 h and visualized by 4xExM. While we rarely found inclusion development severely affected **(a, b)**, we frequently observed inclusions that contain single enlarged particles **(c)** or inclusions that seem to be not affected by the molecule **(d)**. White arrows mark enlarged particles. Scale bar: 8 µm (with 4-fold expansion factor ~2 µm).
